# The Pattern and Staging of Brain Atrophy in Spinocerebellar Ataxia Type 2 (SCA2): MRI Volumetrics from ENIGMA-Ataxia

**DOI:** 10.1101/2024.09.16.613281

**Authors:** Jason W. Robertson, Isaac Adanyeguh, Benjamin Bender, Sylvia Boesch, Arturo Brunetti, Sirio Cocozza, Léo Coutinho, Andreas Deistung, Stefano Diciotti, Imis Dogan, Alexandra Durr, Juan Fernandez-Ruiz, Sophia L. Göricke, Marina Grisoli, Shuo Han, Caterina Mariotti, Chiara Marzi, Mario Mascalchi, Fanny Mochel, Wolfgang Nachbauer, Lorenzo Nanetti, Anna Nigri, Sergio E. Ono, Chiadi U. Onyike, Jerry L. Prince, Kathrin Reetz, Sandro Romanzetti, Francesco Saccà, Matthis Synofzik, Hélio A. Ghizoni Teive, Sophia I. Thomopoulos, Paul M. Thompson, Dagmar Timmann, Sarah H. Ying, Ian H. Harding, Carlos R. Hernandez-Castillo

**Affiliations:** Faculty of Computer Science, Dalhousie University, Halifax, Canada; Sorbonne Université, Institut du Cerveau, INSERM, CNRS, AP-HP, Paris, France; Department of Diagnostic and Interventional Neuroradiology, University Hospital Tübingen, Tübingen, Germany; Department of Neurology, Medical University of Innsbruck, Innsbruck, Austria; Department of Advanced Biomedical Sciences, University of Naples “Federico II”, Naples, Italy; Post-Graduate Program of Internal Medicine, Internal Medicine Department, Hospital de Clínicas, Federal University of Paraná, Curitiba, Brazil; University Clinic and Outpatient Clinic for Radiology, Department for Radiation Medicine, University Hospital Halle (Saale), University Medicine Halle, Halle (Saale), Germany; Department of Electrical, Electronic, and Information Engineering “Guglielmo Marconi”, University of Bologna, Bologna, Italy; Department of Neurology, RWTH Aachen University, Aachen, Germany; JARA-BRAIN Institute Molecular Neuroscience and Neuroimaging, Research Center Jülich GmbH, Jülich, Germany; AP-HP, Hôpital Pitié-Salpêtrière, DMU BioGeM, Department of Genetics, Paris, France; Neuropsychology Laboratory, Department of Physiology, Faculty of Medicine, National Autonomous University of Mexico, Mexico; Center for Translational Neuro- and Behavioral Sciences (C-TNBS), Essen University Hospital, University of Duisburg-Essen, Essen, Germany; Institute of Diagnostic and Interventional Radiology and Neuroradiology, Essen University Hospital, University of Duisburg-Essen, Essen, Germany; Department of Neuroradiology, Fondazione IRCCS Istituto Neurologico Carlo Besta, Milan, Italy; Department of Biomedical Engineering, Johns Hopkins University, Baltimore, USA; Unit of Medical Genetics and Neurogenetics, Fondazione IRCCS Istituto Neurologico Carlo Besta Milan, Italy; Department of Statistics, Computer Science, and Applications “Giuseppe Parenti”, University of Florence, Florence, Italy; Department of Clinical and Experimental Biomedical Sciences “Mario Serio”, University of Florence, Florence, Italy; Clínica DAPI - Diagnóstico Avançado Por Imagem, Curitiba, Brazil; Department of Psychiatry and Behavioral Sciences, Johns Hopkins University, Baltimore, USA; Department of Electrical and Computer Engineering, Johns Hopkins University, Baltimore, USA; Department of Neuroscience and Reproductive and Odontostomatological Sciences, University of Naples “Federico II”, Naples, Italy; Department of Neurodegenerative Diseases, Hertie Institute for Clinical Brain Research, Tübingen, Germany; German Center for Neurodegenerative Diseases (DZNE), Tübingen, Germany; Movement Disorders Unit, Neurology Service, Internal Medicine Department, Hospital de Clínicas, Federal University of Paraná, Curitiba, Brazil; Imaging Genetics Center, Mark and Mary Stevens Institute for Neuroimaging and Informatics, Keck School of Medicine, University of Southern California, Marina del Rey, USA; Department of Radiology, Johns Hopkins University, Baltimore, USA; QIMR Berghofer Medical Research Institute, Brisbane, Australia; School of Translational Medicine, Monash University, Melbourne, Australia

## Abstract

**Objective:** Spinocerebellar ataxia type 2 (SCA2) is a rare, inherited neurodegenerative disease characterised by progressive deterioration in both motor coordination and cognitive function. Atrophy of the cerebellum, brainstem, and spinal cord are core features of SCA2, however the evolution and pattern of whole-brain atrophy in SCA2 remain unclear. We undertook a multi-site, structural magnetic resonance imaging (MRI) study to comprehensively characterize the neurodegeneration profile of SCA2.

**Methods:** Voxel-based morphometry analyses of 110 participants with SCA2 and 128 controls were undertaken to assess groupwise differences in whole-brain volume. Correlations with clinical severity and genotype, and cross-sectional profiling of atrophy patterns at different disease stages, were also performed.

**Results:** Atrophy in SCA2 relative to controls was greatest (Cohen’s *d*>2.5) in the cerebellar white matter (WM), middle cerebellar peduncle, pons, and corticospinal tract. Very large effects (*d*>1.5) were also evident in the superior cerebellar, inferior cerebellar, and cerebral peduncles. In cerebellar grey matter (GM), large effects (*d*>0.8) mapped to areas related to both motor coordination and cognitive tasks. Strong correlations (|*r*|>0.4) between volume and disease severity largely mirrored these groupwise outcomes. Stratification by disease severity showed a degeneration pattern beginning in cerebellar and pontine WM in pre-clinical subjects; spreading to the cerebellar GM and cerebro-cerebellar/corticospinal WM tracts; then finally involving the thalamus, striatum, and cortex in severe stages.

**Interpretation:** The magnitude and pattern of brain atrophy evolves over the course of SCA2, with widespread, non-uniform involvement across the brainstem, cerebellar tracts, and cerebellar cortex; and late involvement of the cerebral cortex and striatum.

## Introduction

Spinocerebellar ataxia type 2 (SCA2) is a rare, inherited neurodegenerative disorder defined by progressive ataxia, dysarthria, dysmetria, sleep disturbances, cognitive dysfunction, slow saccades, peripheral neuropathy, and Parkinsonian signs.^1–3^ It is caused by a CAG triplet expansion in the ATXN2 gene on the long arm of Chromosome 12, which results in a misfolded ataxin-2 protein and toxic gain of function.^1^ This, in turn, results in a neurodegenerative effect that includes Purkinje cell death in the cerebellar cortex; atrophy of the pons, medulla, and cranial nerves; and loss of function in the somatomotor, somatosensory, and vestibular systems, among others.^1^

Magnetic resonance imaging (MRI) approaches have been used to non-invasively describe and quantify structural brain changes in cohorts of individuals with SCA2. For example, voxel-based morphometry (VBM) studies have reported atrophy in the inferior olive, pons, basal ganglia, and corpus callosum, as well as substantial areas of the cerebellar grey and white matter.^3–6^ Diffusion MRI methodologies, which focus on white matter (WM), have found substantial degradation in the cerebellar peduncles, pontine crossing tracts, corticospinal tract, corona radiata, internal capsule, posterior thalamic radiation, medial lemniscus, and corpus callosum.^5,7^ The specific combination of atrophy in the inferior olive, pons, and cerebellum is a telltale signature for SCA2^8^: atrophy levels in these areas plus the dentate nucleus can separate SCA2 from other SCA types using machine learning.^3^

Existing studies of SCA2 brain atrophy have relied on modest sample sizes due to the rarity of the disorder, limiting the statistical power and generalizability of findings. It also remains unclear how the magnitude and profile of brain atrophy evolve with disease progression, and how other clinical variables map onto regional brain atrophy. In this study, we aimed to address these questions using a large, multisite dataset available from the ENIGMA-Ataxia consortium to provide a robust and comprehensive profile of neurodegeneration in people with SCA2, from the pre-symptomatic to severe stages.

## Methods

### Participants and Data

This study used a cross-sectional design with multi-site aggregation through the ENIGMA-Ataxia working group with approval. Demographic, clinical, and imaging data were collected from 110 participants with SCA2 and 128 age- and sex-matched controls (CONT) from eleven different sites: Aachen, Germany; Baltimore, USA; Curitiba, Brazil; Essen/Halle, Germany; Florence, Italy; Innsbruck, Austria; Mexico City, Mexico; Milan, Italy; Naples, Italy; Paris, France; and Tübingen, Germany. The SCA2 and CONT cohorts were age- and sex-matched within each site, and in aggregate (Table 1) showed no significant differences in either sex (*χ*^2^ = 0.0397; *p* = 0.898) or age (Wilcoxon *W* = 7244.5, *p* = 0.700). All data were collected with the participants’ informed consent and human research ethics approval at each contributing site, and were anonymized and assigned new subject identifier codes prior to aggregation. Multi-site processing was governed by the Monash University Human Research and Ethics Committee and the Dalhousie University Research Ethics Board.

**Table 1.** Summary of participant demographics and clinical characteristics.

|  | SCA2 |  |  |  |  |  |  | CONT |  |  |
| --- | --- | --- | --- | --- | --- | --- | --- | --- | --- | --- |
| Site | N | F (%) | Age | Disease Duration | CAG Repeats | SARA | Pre-Symptomatic (%) | N | F (%) | Age |
| Aachen | 2 | 1 (50.0) | 44.5 (24.7) | 4.5 (4.9) | 38.0 (2.8) | 6.3 (1.1) | 0 (0.0) | 3 | 1 (33.3) | 57.7 (7.8) |
| Baltimore | 12 | 9 (75.0) | 49.5 (9.5) | 11.9 (7.8) | 38.0 (1.2) | 12.4 (5.7) | 1 (8.3) | 13 | 11 (84.6) | 51.0 (6.5) |
| Curitiba | 19 | 11 (57.9) | 41.7 (10.9) | 9.4 (7.4) | --<sup>a</sup> | 7.5 (8.3) | 7 (36.8) | 19 | 11 (57.9) | 41.5 (10.9) |
| Essen/Halle | 5 | 2 (40.0) | 49.8 (12.7) | 9.4 (4.7) | 38.6 (1.9) | 15.0 (4.8) | 0 (0.0) | 12 | 4 (33.3) | 51.5 (9.5) |
| Florence | 12 | 3 (25.0) | 45.8 (12.6) | 10.0 (8.0) | 40.5 (1.4) | 12.9 (8.6)<sup>b</sup> | 2 (16.7) | 15 | 7 (46.7) | 49.3 (19.0) |
| Innsbruck | 3 | 1 (33.3) | 39.0 (10.4) | 12.0 (6.2) | 41.7 (3.8) | 17.5 (8.0) | 0 (0.0) | 3 | 2 (66.7) | 37.7 (12.7) |
| Mexico | 13 | 7 (53.8) | 38.5 (15.8) | 11.3 (12.2) | 43.2 (4.8) | 17.0 (7.9) | 0 (0.0) | 15 | 7 (46.7) | 41.7 (13.3) |
| Milan | 15 | 12 (80.0) | 46.3 (9.6) | 11.7 (5.3) | 40.3 (3.1) | 13.6 (7.0) | 0 (0.0) | 20 | 11 (55.0) | 32.8 (6.8) |
| *Milan A* | 2 | 1 (50.0) | 33.5 (0.71) | 4.5 (3.5) | 40.0 (0.0) | 5.8 (2.5) | 0 (0.0) | 6 | 5 (83.3) | 37.5 (9.1) |
| *Milan B* | 13 | 11 (84.6) | 48.2 (8.7) | 12.8 (4.7) | 40.3 (3.3) | 14.8 (6.7) | 0 (0.0) | 14 | 6 (42.9) | 30.8 (4.6) |
| Naples | 13 | 6 (46.2) | 39.9 (11.4) | 8.9 (4.5) | 42.5 (5.5) | 15.0 (4.2) | 0 (0.0) | 13 | 6 (46.2) | 39.2 (11.3) |
| Paris | 12 | 5 (41.7) | 45.0 (12.8) | 9.8 (6.1) | 39.6 (2.9) | 12.6 (6.0) | 0 (0.0) | 12 | 6 (50.0) | 45.0 (12.0) |
| Tübingen | 4 | 2 (50.0) | 33.8 (9.6) | --<sup>c</sup> | 36.0 (2.8) | 1.4 (1.1) | 4 (100.0) | 3 | 1 (33.3) | 30.0 (7.0) |
| **Total** | **110** | **59 (53.6)** | **43.4 (12.1)** | **10.3 (7.3)** | **40.4 (3.8)** | **12.4 (7.6)** | **14 (12.7)** | **128** | **67 (52.3)** | **43.1 (13.0)** |

Parentheses indicate standard deviation except where indicated. Age and disease duration are measured in years. “Milan A” and “Milan B” denote subjects scanned with two different protocols. N: number of subjects; F: number of female subjects; SARA: Scale for the Rating and Assessment of Ataxia score.
<sup>a</sup> Data not available
<sup>b</sup> Converted from equivalent ICARS score<sup>11</sup>
<sup>c</sup> Not applicable; all subjects were pre-symptomatic

Participants with SCA2 were included based on: (1) at least 32 CAG repeats in one allele of the ATXN2 gene^2^, including individuals without clinically manifest ataxic symptoms, and/or (2) genetically confirmed family history of SCA2 and clinical manifestation of disease-consistent ataxia. Age at disease onset was defined as the point at which the participants first reported or were recorded clinically as showing ataxic symptoms. Ataxia severity at the time of MRI collection was determined using the Scale for Assessment and Rating of Ataxia (SARA; 10 sites)^9^ and/or the International Cooperative Ataxia Rating Scale (ICARS; 3 sites).^10^ For analytical consistency, the data from the one site that only collected ICARS scores (Florence) were converted to SARA based on the linear regression reported by Rummey et al.^11^ To ensure the validity of this approach, converted scores were also calculated for participants from the other two ICARS sites (Baltimore and Essen/Halle), for whom scores on both scales are available; no significant difference was found between the calculated and collected SARA scores (Supplementary Table S1).

A whole-brain, high-resolution T1-weighted structural MRI image was collected for each subject. Imaging data collection protocols varied by site, but were consistent across patients and controls. Scanners had a field strength of 3 T (10 sites) or 1.5 T (1 site), and were manufactured by either Siemens (Erlangen, Germany; 7 sites) or Phillips (Amsterdam, Netherlands; 4 sites). One site (Milan) collected protocols on two different scanners, and was thus treated as two different sites for the purpose of analysis. Full imaging protocols are summarized in Supplementary Table S2.

### Image Processing

All T1-weighted images were pre-processed using a previously described pipeline for the ENIGMA-Ataxia project^12^, which combines specialized software for analyzing the cerebellum and cortex.^13^ First, a cerebellar mask was derived for each image then visually inspected. Minor segmentation errors could be corrected manually, while more substantial errors resulted in exclusion. This step could be performed entirely manually, or automated using the ACAPULCO^14^ algorithm (v0.2.1). Next, cerebellar grey matter (GM) was extracted using the Spatially Unbiased Infratentorial Toolbox (SUIT) v3.2 toolbox^15^ for SPM12 v7771^16^, as implemented for MATLAB (Natick, MA). Using the previously generated cerebellar mask, the images were segmented into GM and WM partial volumes, which were DARTEL normalized and resliced into SUIT space; Jacobian modulation was used to ensure that each voxel remained proportional to its original volume. The resulting cerebellar GM images were inspected for normalization errors, and the spatial covariance of these images was calculated and compared across the entire cohort: outliers were visually inspected to determine if rejection was necessary due to processing errors. Finally, the images were spatially smoothed using a Gaussian kernel with a full width at half-maximum (FWHM) of 3 mm.

As SUIT is designed specifically for cerebellar analysis, whole-brain WM and cerebral GM were instead quantified using the Computational Anatomy Toolbox (CAT12, v12.5)^17^ for SPM12. Images were bias-field corrected and skull stripped, then segmented into GM, WM, and cerebrospinal fluid (CSF) partial volumes from which the total intracranial volume (ICV) could be calculated. The GM and WM partial volumes were then DARTEL registered to Montreal Neurological Institute (MNI) standard space with Jacobian modulation to maintain volume encoding. The GM image was masked to remove the cerebellum, so its data would not influence cerebral GM findings. Both GM and WM images were then smoothed with a 5 mm FWHM Gaussian kernel.

### Statistical Analysis

Statistical analyses were performed using SPM12 in MATLAB. In all cases, heteroscedasticity was assumed and corrected for, with final voxel-level inferences estimated with family-wise error (FWE) corrections at the *α* = 0.05 level, using random field theory to account for the large number of multiple comparisons endemic to imaging analysis^18^.

#### Between-Group Analysis for Disease Effects

Groupwise comparisons between SCA2 and CONT participants were performed by creating a generalized linear model in SPM12 with Group (2 levels) and Site (12 levels, treating Milan as two sites) as factors and ICV and Age as covariates. Of these, only Group is a comparison of interest; the remainder are included as nuisance variables.

Inferences were first undertaken at the voxel level, then for interpretative purposes the results were mapped to the following anatomical atlases: the Harvard-Oxford cortical and subcortical GM atlases and the Johns Hopkins University cortical WM atlas as included with FSL^19^; the SUIT cerebellar grey matter and dentate atlases^20,21^; the van Baarsen cerebellar WM atlas^22^; and the FreeSurfer brainstem atlas.^23^ Additionally, functional correlates were mapped using the multi-domain task-based (MDTB) parcellation developed by King et al.^24^

#### Clinical Correlations for SCA2 Patients

The effects of healthy aging^25^ and site-specific confounds^26^ were estimated and adjusted for prior to undertaking clinical correlations using previously reported methods.^27^ These adjustments are described in detail in the Supplemental Methods.

The age- and site-adjusted SCA2 images were assessed for linear relationships between GM or WM volume and three clinical measures: number of CAG repeats, disease duration, and SARA score. For each of these correlations, ICV was again included as a nuisance covariate to account for the effect of subject head size.

#### Disease Staging

Cross-sectional analyses were performed on subsets of the SCA2 participant data to better understand how the disease progresses. The data were subdivided into five groups – pre-symptomatic (SARA < 3)^28^ and four quartiles of symptomatic individuals (quartile separation values: SARA = 9, 13.5, and 18) – which were then each compared to the full CONT cohort. Because SCA2 subject counts were necessarily much smaller than the full dataset, the age- and site-corrected data were calculated then compared using a generalized linear model with Group (2 levels) as the only factor and ICV as the only nuisance covariate. Voxel-level inferences were calculated using FWE as described above.

An additional categorical split was undertaken by dividing the SCA2 cohort into four subgroups based whether participants had above- or below-median CAG repeat length (median: 40 repeats) and/or disease duration (median: 9 y). Groupwise analyses were performed between each subgroup and the full CONT cohort using age- and site-corrected data as described above.

#### Effect Size

Once all statistics were calculated and thresholded at *p*_FWE_ < 0.05, the resulting map of T-values was converted voxelwise to Cohen’s *d* for groupwise comparisons, or Pearson’s *r* for correlations, to express effect size. The details of this are described in the Supplemental Methods. The converted images of statistically significant *d* and *r* values were used for all further data reporting and visualization.

## Results

### Volumetric Differences Between SCA2 Patients and Healthy Controls

Voxel-level statistical differences between SCA2 and CONT groups are presented in Figure 1. The strongest effects were observed in WM regions (Figure 1A), with the greatest atrophy (*d* > 1.5) observed in all cerebellar and cerebral peduncles, the pons, the cerebellar subcortex, and the corticospinal tract bilaterally. More moderate atrophy was also observed in many areas of the cerebral WM, including the internal capsule, superior fronto-occipital fasciculus, and superior corona radiata, particularly in areas encompassing motor tracts. Parts of the body and splenium of the corpus callosum also showed significant atrophy in SCA2 compared to CONT.

**Figure 1.**
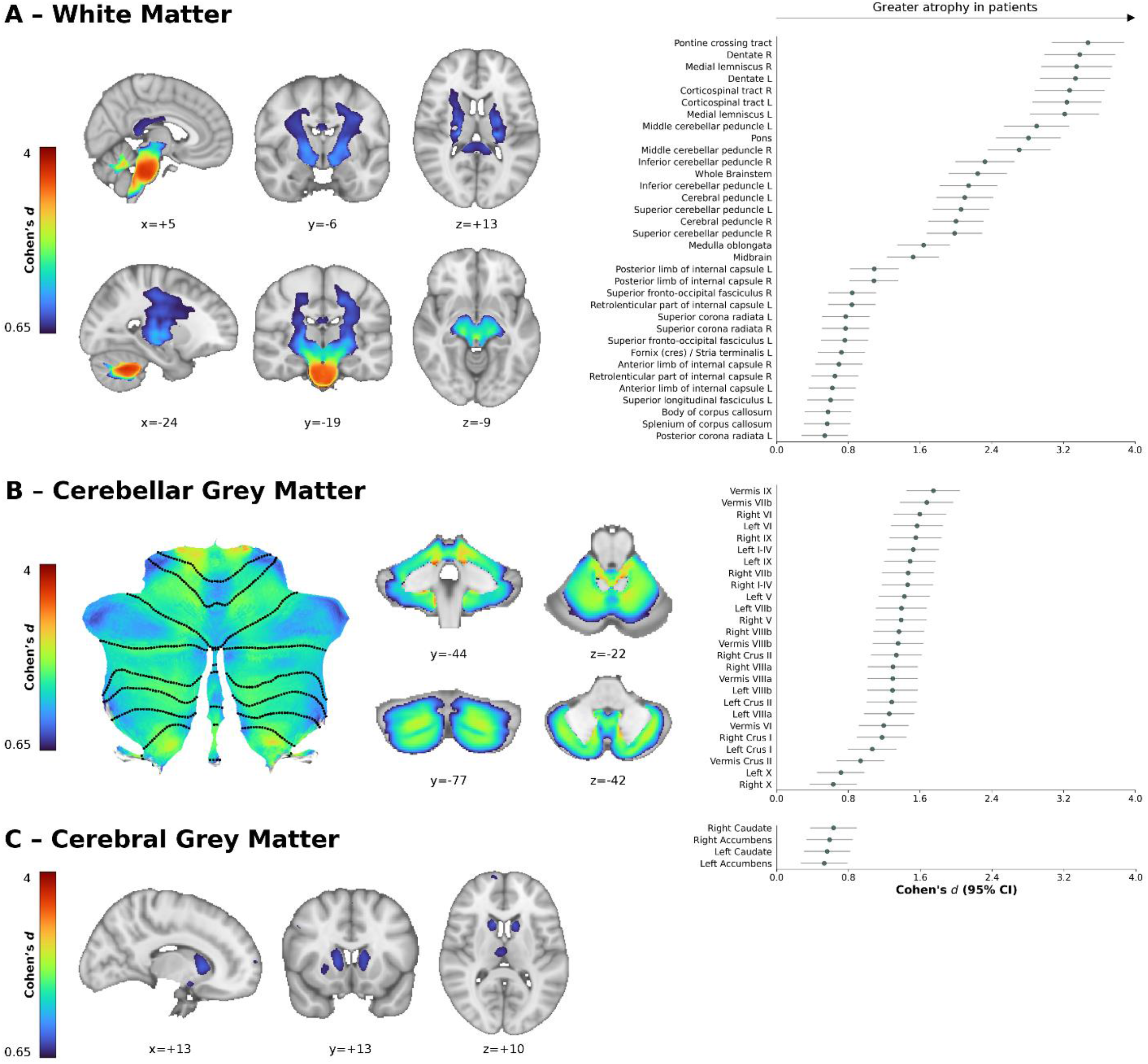
Regions of significantly lower volume (voxel-level FWE-corrected *p* < 0.05) in participants with SCA2 relative to CONT in the (A) whole brain white matter, (B) cerebellar grey matter, and (C) cerebral grey matter. Left: representative slices or cerebellar flatmaps illustrating the areas of significant atrophy in SCA2 subjects. Right: forest plots illustrating regional effects (Cohen’s *d* > 0.5, reflecting moderate or greater effect size); error bars represent the 95% confidence interval (CI). Slice coordinates are in Montreal Neurological Institute (MNI) space.

With respect to the cerebellar GM (Figure 1B), significant volume loss was observed across the entirety of the cerebellum, with all anatomical regions except Lobule X showing a large (*d* > 0.8) bilateral atrophic effect. Areas of relatively greater atrophy were seen in bilateral lobules I-IV and IX.

Finally, considering the cerebral GM (Figure 1C), significant atrophy was restricted to moderate effect sizes (*d* > 0.5) in the bilateral caudate nucleus, amygdala, nucleus accumbens, and thalamus.

### Network Representation of Patient-Control Differences

We subsequently mapped the cerebellar GM atrophy profiles (Figure 2A) to the functional regions of the cerebellum defined by the multi-domain task battery (MDTB) atlas (Figure 2B).^24^ Atrophy was most pronounced in MDTB regions 5-7 – associated with attention, executive function, and language/emotion processing – followed by MDTB regions 1, 2, and 4, which are primarily related to motor generation and planning.

**Figure 2.**
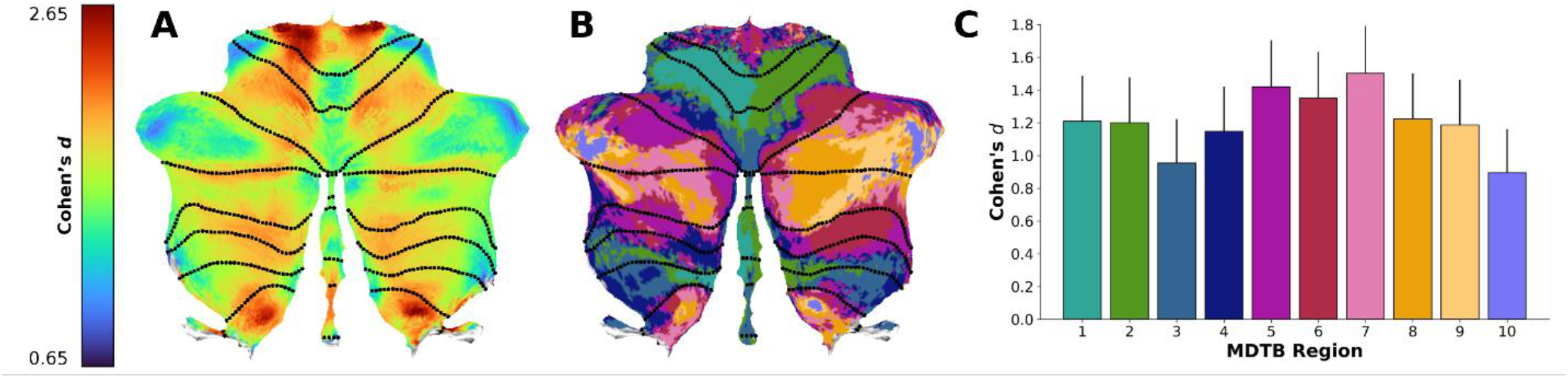
Mapping of structural changes to functional networks in the cerebellum. (A) Cerebellar flatmap of the voxel-level effect size of volume differences in participants with SCA2 vs. CONT in cerebellar grey matter, recalled from Figure 1B. (B) Cerebellar flatmap of the multi-domain task battery (MDTB) functional atlas from King et al.^24^ (C) A bar chart of the effect sizes by MDTB region; bar height is the mean effect size within each region, while error bars are the positive 95% CI. The colours of the bars in Panel C match the colours of the regions in Panel B.

### Clinical Correlations in SCA2 Patients

As illustrated in Figure 3A, SARA scores were strongly inversely correlated with WM volume in the cerebellar peduncles and pons (|*r*| > 0.6), with more moderate correlations in the cerebrum (|*r*| > 0.4), specifically in the anterior portion of the left superior longitudinal fasciculus and regions of the bilateral premotor and supplementary motor tracts. In the GM (Figure 3B), significant negative correlations with SARA were observed throughout much, but not all, of the cerebellum. The areas with strongest correlations were bilateral lobules I-IV, VIIIa/b, IX, and crus II. In the cerebrum, significant negative correlations with SARA were observed in the left caudate nucleus, inferior and middle frontal gyri, and precentral gyri.

**Figure 3.**
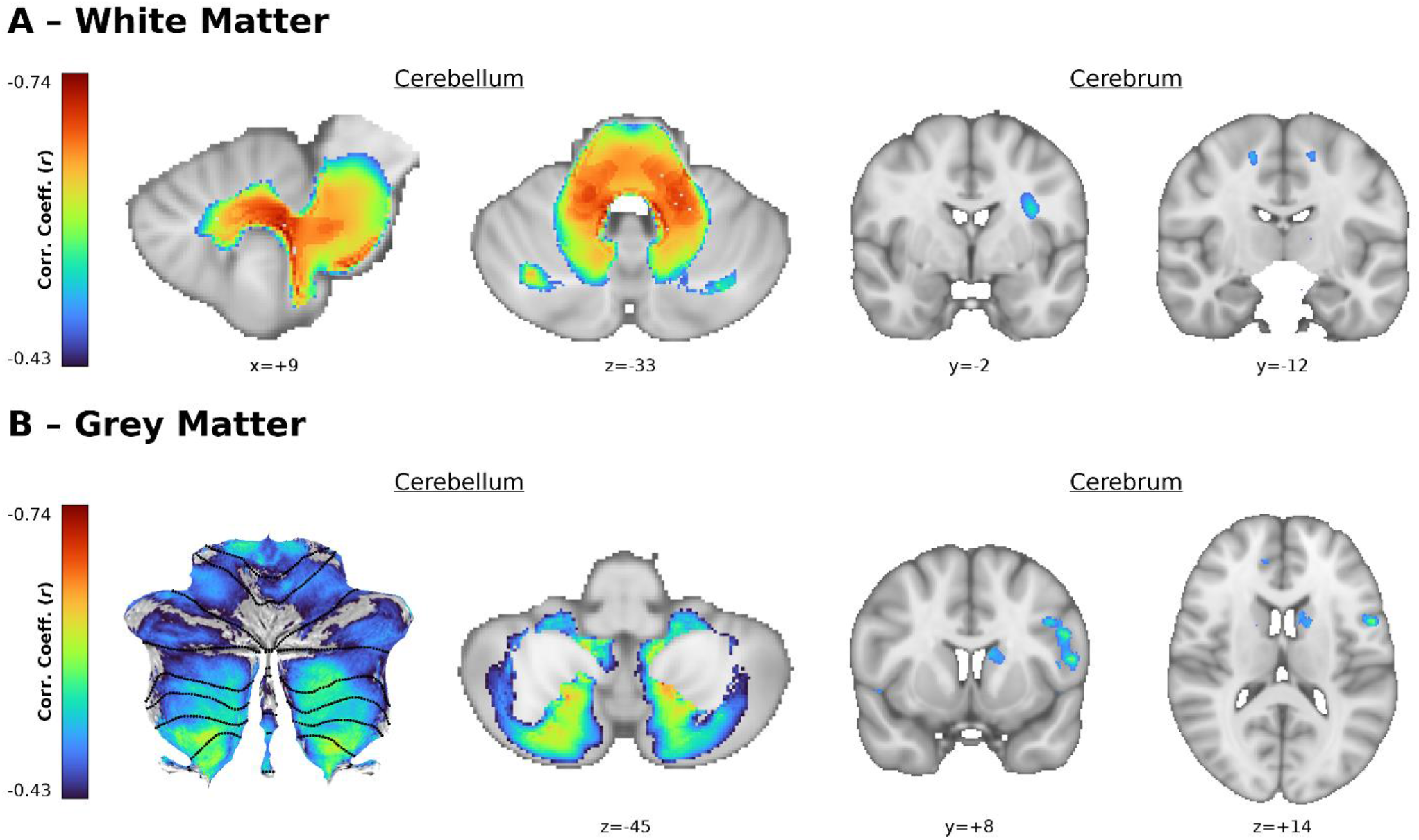
Correlations between regional volume and ataxia severity (SARA score) in SCA2 participants (voxel-level FWE-corrected *p* < 0.05). (A) SARA vs. white matter volume in the cerebellum and brainstem (left) and cerebrum (right). (B) SARA vs. grey matter volume in the cerebellum (left) and cerebrum (right). Slice coordinates are in MNI space.

Some localized correlations were also observed with CAG repeats in the cerebral WM, in the area of the left superior longitudinal fasciculus and superior corona radiata (Supplemental Figure S1). No significant correlation was found between disease duration and any volumetric measure.

### Disease Staging

Cross-sectional stratification of the SCA2 cohort by SARA score is presented in Figure 4. As a broad trend, tissue atrophy became more severe and widespread as disease severity increased: whereas pre-symptomatic participants showed only isolated atrophy in the cerebellar GM and WM, participants in the third or fourth quartiles of SARA score showed substantial atrophy across the same regions as seen in the full analysis (*cf*. Figure 1). Interestingly, whereas pre-symptomatic, first- and second-quartile participants showed no significant atrophy in the cerebral GM at all, significant volume loss was evident in areas of the motor and premotor cortices in participants in the fourth quartile of SARA score.

**Figure 4.**
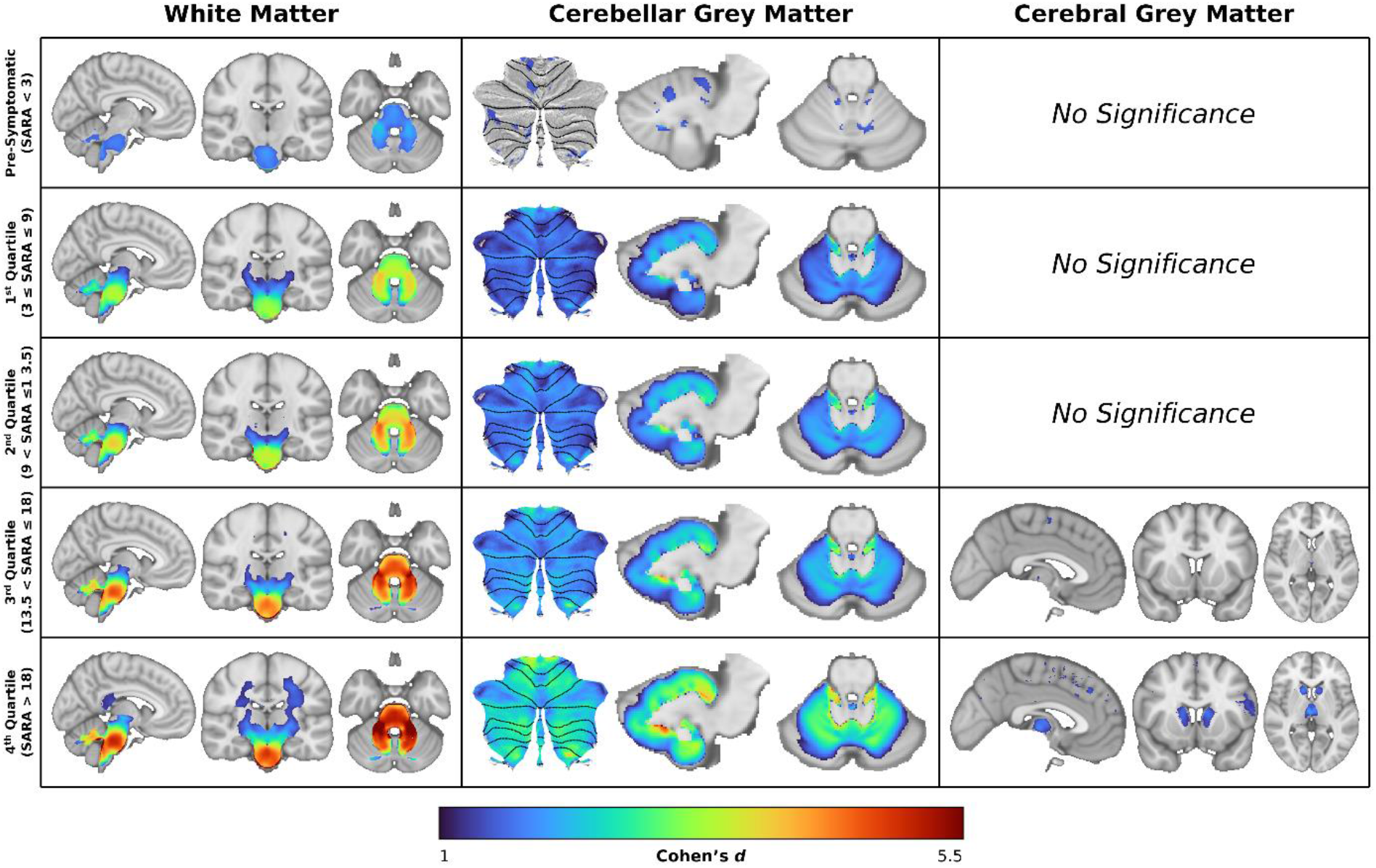
Stratification of the SCA2 cohort based on ataxia severity (SARA score). Voxel-level effect size maps (FWE-corrected *p* < 0.05) of each SCA2 subgroup relative to CONT. From top to bottom: Pre-Symptomatic (SARA < 3; *n* = 14); 1^st^ Quartile (3 *≤* SARA *≤* 9; *n* = 26); 2^nd^ Quartile (9 < SARA *≤* 13.5; *n* = 22); 3^rd^ Quartile (13.5 < SARA *≤* 18; *n* = 23); 4^th^ Quartile (SARA > 18; *n* = 24). All images show representative slices or flatmaps of the affected tissue type: white matter (left) is at MNI coordinates (−6, -20, -30); cerebellar grey matter (centre) shows the flatmap and slices from MNI coordinates *x* = -10 and *z* = -25; cerebral grey matter (right) is at MNI coordinates (−2, +11, +4).

Cross-sectional stratification based on CAG repeat length and disease duration is presented in Figure 5. In participants with low CAG repeats, there were no significant results in the cerebral GM regardless of disease duration, whereas the magnitude of atrophy in the cerebellar GM was modulated by disease duration. In participants with high CAG repeats, a greater magnitude and spatial extent of atrophy was observed relative to those in the low CAG group, with a similar disease duration-related trend. Separate analysis was run to determine if there was any interaction effect between CAG repeat length and disease duration beyond the additive results shown here, however no significant interaction was found.

**Figure 5.**
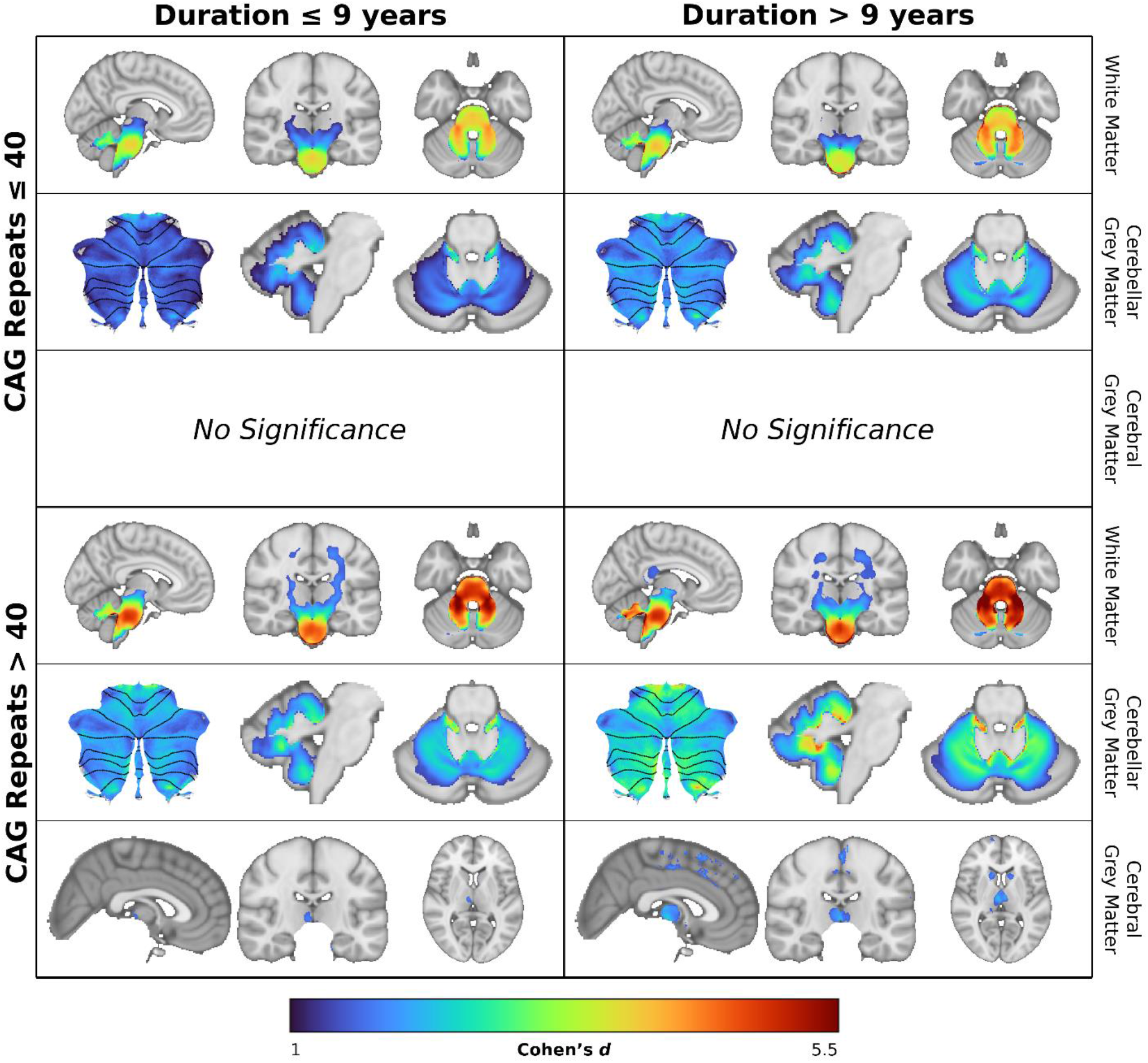
Stratification of the SCA2 cohort based on CAG repeat length and disease duration. Voxel-level effect size maps (FWE-corrected *p* < 0.05) of each SCA2 subgroup relative to healthy controls. *Top left*: CAG repeats and disease duration below the median of each measure (CAG repeats *≤* 40; Duration *≤* 9 y; *n* = 26). *Top right*: CAG repeats below the median and disease duration above the median (CAG repeats *≤* 40; Duration > 9 y; *n* = 21). *Bottom left*: CAG repeats above the median and disease duration below the median (CAG repeats > 40; Duration *≤* 9 y; *n* = 15). *Bottom right*: CAG repeats and disease duration above the median of each measure (CAG repeats > 40; Duration > 9 y; *n* = 16). Within each cell, representative images appear from the whole-brain white matter (top) at MNI coordinates (−6, -20, -30); the cerebellar grey matter (middle) as the flatmap and slices from MNI coordinates *x* = -5 and *z* = -26; and cerebral grey matter (bottom) at MNI coordinates (+2, -18, +4).

## Discussion

In this large, multi-site, quantitative MRI study of participants with SCA2 and matched controls, we provide a robust assessment of the pattern and evolution of brain atrophy in the largest cohort of SCA2 carriers collected to date. We observed the greatest atrophy in the cerebellar WM and pons, followed by smaller but still substantial effects in the cerebellar GM and cerebral WM pathways between the cerebellum and the motor cortex, and moderate effects in areas of the basal ganglia related to motor coordination. This pattern holds true in the main groupwise comparisons, when examining correlations with disease severity, and when examining different stages of disease progression. Cerebellar functional mapping shows atrophy in regions related to both motor and cognitive functions.

### The Pre-Eminence of White Matter Atrophy in SCA2

Atrophy was greatest in the WM of the cerebellum, pons, and corticospinal tracts, consistent with the overall characterization of SCA2 as an olivopontocerebellar disease.^8^ The fact that the WM effect size is so much greater than that of any GM results, combined with the extensive WM atrophy observed in pre-symptomatic participants, clearly implicates WM atrophy as the first observable component of SCA2 neurodegeneration. This finding is consistent with previous neurochemical research in both humans and murine models, which has suggested that dysfunctional lipid synthesis leading to demyelination plays an important role in the pathophysiology of SCA2.^29–31^

Our findings corroborate and extend previous reports that brain atrophy precedes the onset of ataxia symptoms in individuals with SCA2.^6,28,32^ These effects were particularly evident in the pons and cerebellar WM, as well as the anterior and superior posterior lobes of the cerebellar GM. This is in line with Jacobi et al.^28^, who also found significant WM loss in the cerebellum and pons compared to controls, as well as GM loss in Lobules V and VI, despite the participants not being projected to show clinical signs of ataxia for about a decade. Nigri and colleagues^6^ similarly report reduced pons volume in pre-symptomatic SCA2 carriers relative to controls.

This preponderance of abnormalities in WM extends to our cross-sectional analysis of symptomatic SCA2 participants: stratifying the SCA2 cohort based on disease severity produces a pattern of increasing atrophy magnitude within the affected areas at each stage, as well as a progressive expansion of affected regions. Irrespective of disease stage, however, the atrophy pattern is similar: maximal in the pons and cerebellar WM, followed by the cerebellar GM, then finally the cerebral subcortical and cortical regions. These cross-sectional findings agree with previously reported longitudinal changes in SCA2, which tend to show widespread loss of integrity in the WM of SCA2 patients with time, most commonly in the corpus callosum, pons, internal capsule, and cerebellar peduncles^5,6,33–35^. By comparison, fewer longitudinal studies have reported GM data. One early study by Mascalchi et al.^35^ found significant longitudinal atrophy in the GM of the superior vermis and flocculonodular lobules in the cerebellum; and the left midfrontal gyrus, basal ganglia, and thalamus in the cerebrum. Another by Nigri et al.^6^ found longitudinal atrophy in the cerebellum as a whole, as well as lobules V, VI, and VIIA. Taken together, these studies and ours demonstrate an evolutionary pattern in SCA2-related brain atrophy.

### Connecting Clinical and Imaging Data

Ataxia severity was most strongly related to volume in the cerebellar and pontine WM, and the anterior and inferior posterior lobes of the cerebellar GM. Previous work has similarly shown correlations between disease severity and atrophy in the cerebellar and pontine WM^4,5,7,36^, even in pre-symptomatic subjects.^37,38^ Strong correlations were also found in the few studies that examined cerebellar GM: Della Nave et al.^36^ found a negative correlation between ICARS score and cerebellar GM volume, while Jung et al.^4^ found negative correlations between functional scores and age-adjusted volumes in Lobules IV, VIIb, and VIII; and Crus I. The correlations in our data covered a more widespread area of the cerebellar GM than others, however the regions of strongest correlation are consistent with prior findings, suggesting that this may be a reflection of the greater statistical power of our sample. Correlations in cerebral regions were more isolated and do not entirely correspond to previous outcomes^7^, however many of the regions highlighted by the present study — specifically, the inferior frontal, middle frontal, and precentral gyri — can be connected to cognitive and executive functions such as language processing and performance^39–42^, risk analysis^43^, and memory^39^, more so than motor control. Thus, these areas warrant further investigation with respect to the cognitive components of SCA2.

By contrast, both the literature and present data are ambiguous regarding the relationship between CAG repeats or disease duration and neuroimaging markers. While we show clear atrophic effects when comparing data above vs. below the median of each measure, only a small, localized correlation exists between CAG repeats and neural volume, and none with disease duration. Prior studies also report varied relationships: while some show strong negative correlations between disease duration and volume in the cerebellum, pons, and medulla, among other areas^4,6^, others find none.^34,36^ Similarly, Reetz et al.^38^ found negative correlations between CAG repeats and volume in the pons, cerebellum, and whole brainstem, while Tu et al.^34^ did not. These discrepancies may be because most studies capture a single snapshot in time, whereas these clinical variables may relate more to the *rate* of degeneration: the clearer evidence of relationships between CAG repeats, disease duration, and neurodegeneration in the staging analysis lends some credence to this notion, however future longitudinal studies would be necessary to confirm it.

### Functional Correlates of Cerebellar GM Atrophy

Cerebellar regions tend to be described in terms of motor or non-motor processes, however this categorization has not always been consistent: Hernandez-Castillo et al.^44^ observed that, depending on the cerebral region a given lobe was functionally connected to, it could relate to motor or non-motor outcomes. Olivito et al.^45^ subsequently found, among other results, reduced functional connectivity between Lobule/Vermis VI and cortical regions associated with both motor and non-motor functionality in participants with SCA2.

Recent efforts to functionally map the cerebellum without regard for cerebellar lobular boundaries, such as that by King et al.^24^, have enabled more nuanced descriptions. Using this map, we found large effect sizes in regions related to both motor and non-motor processes, with slightly higher effect sizes in regions related to cognitive processes, particularly in the domains of attention, working memory, and language and emotional processing. Deficits in these domains are concordant with cerebellar cognitive affective syndrome, an important component of SCA2.^46^ These same domains are also affected by atrophy in the cerebral areas found to correlate with SARA, as noted previously. Thus, while the present study did not include cognitive data, the identified atrophy pattern is consistent with the known clinical profile of SCA2.

### Limitations

The contributions of this study are not without limitations. First, while VBM does provide a useful illustration of WM atrophy, diffusion MRI would provide a clearer picture of the microstructural breakdown in SCA2 patients’ WM. This would be particularly powerful at the scale of subject data available to ENIGMA-Ataxia. Second, while our cross-sectional results do generally agree with those of previous longitudinal studies and provide valuable insights into disease expression and progression, the identification and validation of biomarkers for use as surrogate trial outcome measures or clinical tracking will necessitate future large-scale longitudinal assessments. Third, we cannot draw conclusions about how the observed neurodegeneration in SCA2 participants relates to symptoms beyond ataxia severity, such as cognitive decline, because that data is not consistently available across all sites.

Prospective studies with standardized behavioural task batteries are indicated. Finally, our between-sites harmonization relies on well-accepted regression-based approaches, however we acknowledge that more sophisticated methods for addressing between-sites variability in neuroimaging data exist that would be worthy of further investigation.^47^

### Conclusion

In this study, we have comprehensively characterized the pattern and spread of neurodegeneration in SCA2 based on the largest sample size of MRI images to date. We have demonstrated not only that symptom-relevant atrophy in the cerebellum and brainstem continues to progress over time, starting before clinical onset, but also that the cerebral cortex – particularly the motor system – becomes increasingly impacted in later disease stages. We have also shown that the greatest neurodegenerative effects are consistently found in cerebellar WM; followed by the cerebellar GM; then finally the cerebral motor tracts, thalamus, and striatum. Together, these observations paint a picture of a disease spreading outward from its origin point in the cerebellum and brainstem to eventually involve large swaths of the motor system.

## Acknowledgments

This work was supported by the following grants and funding agencies: Natural Sciences and Engineering Research Council (NSERC) grant number RGPIN-2022-03368 and Canada Research Chair (CRC) grant number CRC-2020-00079 to Carlos R. Hernandez-Castillo; German Federal Ministry of Education and Research (BMBF) grants 01GQ1402 and 01DN18022 to Kathrin Reetz; the National Ataxia Foundation Clinical Research Consortium, National Science Foundation (NSF) grant EEC-1460674, National Institute of Health (NIH) National Institute of Neurological Disorders and Stroke (NINDS) grant R01NS056307, Johns Hopkins Institute for Clinical and Translational Research support from NIH National Center for Advancing Translational Sciences (NCATS) grant UL1TR001079 and the Jane Tanger Black Fund for Young-Onset Dementia Research to Chiadi U. Onyike; German Research Foundation (DFG) grants DE 256/1-1 and TI 239/17-1 to Dagmar Timmann; National Council of Science and Technology of Mexico (CONACYT) grant A1-S-10669 and General Directorate of Academic Personnel Affairs Support Program for Research and Technological Innovation Projects (DGAPA-PAPIIT) grant IN214122 to Juan Fernandez-Ruiz; Italian Ministry of Health grant RF 2011-02437420 to Caterina Mariotti; the Italian Association for the Fight Against Ataxic Syndromes (AISA), Campania Section to Sirio Cocozza; DFG grant 4414096270 to Matthis Synofzik; Australian National Health and Medical Research Council Ideas Grant 1184403 and Investigator Grant 2026191 to Ian H. Harding; NIH Big Data to Knowledge (BD2K) award U54 EB020403 and NIH grants R01MH123163, R01MH121246, and R01MH116147 to Paul M. Thompson. For a complete list of ENIGMA-related grant support please see here: http://enigma.ini.usc.edu/about-2/funding/.

The authors would also like to acknowledge the contributions of Victor Gadelha, Hellen Della-Justina, Salmo Raskin, and Arnolfo de Carvalho Neto during data acquisition in Curitiba.

## Author Contributions

JWR performed post-aggregation data analysis and led the writing of the manuscript. IHH and CRH conceived the study, co-ordinated data acquisition, and directed data analysis. PMT founded and directs the ENIGMA consortium and provided important feedback on the manuscript. IA, BB, SB, AB, SC, LC, A. Deistung, SD, A. Durr, SLG, MG, C. Mariotti, C. Marzi, MM, FM, WN, LN, AN, SEO, JLP, KR, SR, DT, and SHY performed imaging data collection at their respective sites. IA, AB, SC, A. Deistung, CRH, and WN performed imaging data analysis at their respective sites. SB, LC, ID, A. Durr, MG, C. Mariotti, MM, FM, WN, LN, CUO, KR, FS, HAGT, DT, and SHY performed clinical data colletion at their respective sites. SB, SC, A. Durr, JF, IHH, CRH, C. Mariotti, MM, FM, CUO, KR, MS, HAGT, and DT served as principal investigators for their respective sites SH and CRH performed analytical methods development for the ENIGMA-Ataxia pipelines. JLP and SIT performed additional analyses.

## Data Availability

All code and data processing instructions are available at https://github.com/HardingLab/enigma-ataxia. The data may be requested for further research purposes through a secondary data use proposal submitted to the ENIGMA-Ataxia working group (http://enigma.ini.usc.edu/ongoing/enigma-ataxia).

