## Supplementary Table for "The Pattern and Staging of Brain Atrophy in Spinocerebellar Ataxia Type 2 (SCA2): MRI Volumetrics from ENIGMA-Ataxia"

### Supplementary Methods

*Age and Site Correction*

Site effects were calculated on a voxelwise basis using CONT subjects as a template, using a method previously described by Harding et al.[1] The mean CONT image for each site $\mu_{site}$ was subtracted from the global mean $\mu_{global}$ to produce a site-correction image; this was then added to each pre-processed subject image $y$ from that site to produce the site-adjusted image $\hat{y}$:

$$\hat{y} = y + (\mu_{global} - \mu_{site})$$

To determine the healthy aging effect, a linear regression was run on all voxels in each CONT image with respect to subject age $x$:

$$y = \beta_{age}x + \beta_0 + \epsilon$$

This regression constant $\beta_{age}$ was then used to calculate and adjust for the effect of subject age $x$ relative to mean age $\bar{x}$ on the SCA2 subjects' site-adjusted images:

$$y_{adj} = \hat{y} - \beta_{age}(x - \bar{x})$$

This final, adjusted image $y_{adj}$ was used for all within-cohort clinical correlations.

*Expressions of Effect Size*

Between-groups comparisons were converted to Cohen's *d* using the formula

$$d = \frac{t(n_1 + n_2)}{\sqrt{n_1 n_2\, df}}$$

where $n_1$ and $n_2$ were the sample sizes of each group, $t$ was the T-score in each voxel, and $df$ represented the degrees of freedom of the comparison.[2] The confidence interval (CI) limits were then calculated as

$$d \pm 1.96\sqrt{\frac{n_1 + n_2}{n_1 n_2} + \frac{d^2}{2(n_1 + n_2)}}$$

based on a previously published formula.[3] Regressions were converted to Pearson's *r* using the formula[2]

$$r = \frac{t}{\sqrt{t^2 + df}}$$

### Supplementary Figures

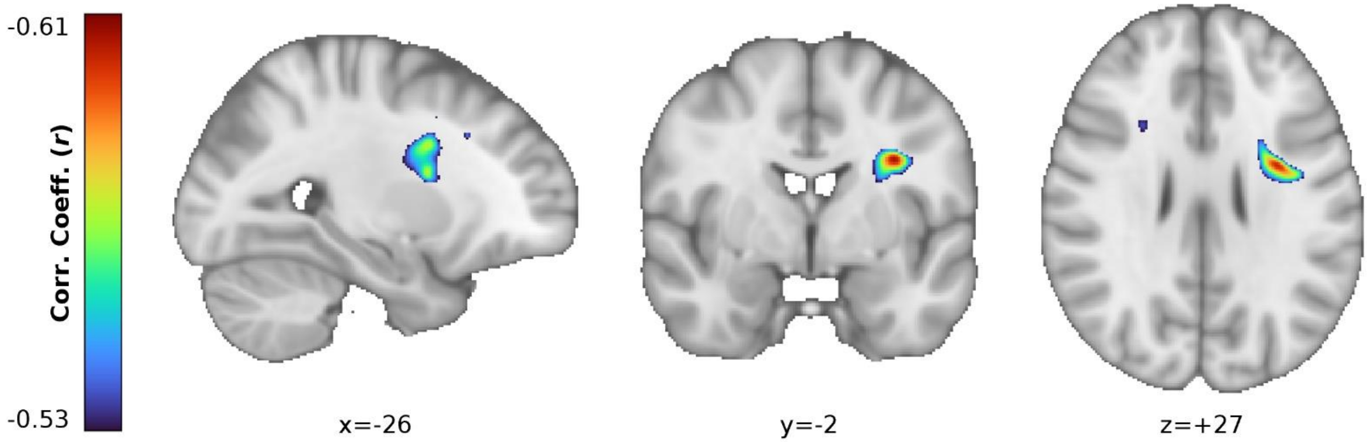


**Supplementary Figure S1.** Representative slices illustrating voxel-level regressions between white matter volume and CAG repeats in age- and site-corrected scans from SCA2 patients, with intracranial volume as a nuisance regressor. Slice coordinates are in MNI space. Note that the colour scale does *not* match that of Figure 3.

### Supplementary Tables

**Supplementary Table S1.** Comparison of converted SARA scores derived from ICARS and SARA scores collected directly from the subjects.

| Subject ID | ICARS | Converted SARA | True SARA |
|---|---|---|---|
| baltimore_sca2_03 | 16 | 6 | 6 |
| baltimore_sca2_06 | 38 | 16 | 16.5 |
| baltimore_sca2_07 | 27 | 11 | 12 |
| baltimore_sca2_10 | 26 | 10.5 | 11 |
| baltimore_sca2_11 | 13 | 4.5 | 2.5 |
| baltimore_sca2_12 | 37 | 15.5 | 15.5 |
| baltimore_sca2_15 | 23 | 9 | 10.5 |
| baltimore_sca2_16 | 53 | 22.5 | 23 |
| baltimore_sca2_17 | 28 | 11.5 | 10.5 |
| baltimore_sca2_18 | 44 | 18.5 | 16 |
| baltimore_sca2_19 | 21 | 8 | 7.5 |
| baltimore_sca2_20 | 42 | 17.5 | 18 |
| essen_sca2_01 | 25 | 10 | 14.5 |
| essen_sca2_02 | 32 | 13 | 13 |
| essen_sca2_03 | 52 | 22 | 22 |
| essen_sca2_04 | 40 | 16.5 | 16.5 |
| essen_sca2_05 | 15 | 5.5 | 9 |

Shapiro-Wilk tests of normality: converted SARA ($W$ = 0.959; $p$ = 0.619); true SARA ($W$ = 0.984; $p$ = 0.987)
Paired-samples t-test results: $t(16)$ = -0.932; $p$ = 0.365
Conversion formula from Rummey et al.[4]

**Supplementary Table S2.** Summary of imaging protocols as reported by each site.

| | Aachen | Baltimore | Curitiba | Essen/Halle | Florence | Innsbruck | Mexico | Milan[a] | Naples | Paris | Tübingen |
|---|---|---|---|---|---|---|---|---|---|---|---|
| **Scanner** | Siemens Prisma | Philips Intera | Siemens Skyra | Siemens Biograph | Philips Intera | Siemens Verio | Philips | Philips Achieva | Siemens Trio | Siemens Trio | Siemens Skyra |
| **Field (T)** | 3 | 3 | 3 | 3 | 1.5 | 3 | 3 | 3 | 3 | 3 | 3 |
| **Channels** | 64 | 8 | 16 | 16 | 6 | 12 | 32 | 32 | 8 | 32 | 32 |
| **Sequence** | MPRAGE | MPRAGE | MPRAGE | MPRAGE | IR-3D GRE | MPRAGE | FFE | TFE | MPRAGE | MPRAGE | MPRAGE |
| **TR (ms)** | 2500 | 10.3136 | 2530 | 2530 | 8.1 | 1800 | 8 | 9781 | 1900 | 2530 | 2300 |
| **TE (ms)** | 4.37 | 6 | 3.36 | 3.26 | 3.7 | 2.18 | 3.7 | 4.6 | 3.4 | 3.65 | 2.32 |
| **TI (ms)** | 1100 | --[b] | 1100 | 1100 | 764 | 900 | 900 | --[b] | 900 | 900 | 900 |
| **Flip Angle (°)** | 7 | 8 | 7 | 7 | 8 | 9 | 25 | 8 | 9 | 9 | 8 |
| **Plane** | Sagittal | Axial | Sagittal | Sagittal | Sagittal | Coronal | Sagittal | Sagittal | Axial | Sagittal | Sagittal |
| **Slices** | 192 | Variable | 176 | 176 | 160 | 160 | 176 | 185 | 160 | 160 | 192 |
| **Field of View** | 256 × 256 | 256 × 256 | 256 × 256 | 256 × 256 | 256 × 256 | 220 × 178 | 256 × 256 | 240 × 240 | 256x192 | 256 × 256 | 230 × 230 |
| **Voxel Size (mm, X × Y × Z)** | 1 × 1 × 1 | 0.828125 × 0.828125 × 1.1 | 1 × 1 × 1 | 1 × 1 × 1 | 1 × 1 × 1 | 0.43 × 0.43 × 1.2 | 1 × 1 × 1 | 1 × 1 × 1 | 1 × 1 × 1 | 1 × 1 × 1 | 0.9 × 0.9 × 0.9 |

FFE: fast field-echo; IR-3D GRE: 3D inversion recovery gradient echo; MPRAGE: magnetization-prepared rapid gradient-echo; TE: echo time; TFE: turbo field-echo; TI: inversion time; TR: repetition time

[a] Protocol only used for a subset of subjects.

[b] Information not available.
